# Developmental Alcohol Exposure in Drosophila: Effects on Adult Phenotypes and Gene Expression in the Brain

**DOI:** 10.1101/2021.04.22.440913

**Authors:** Sneha S. Mokashi, Vijay Shankar, Rebecca A. MacPherson, Rachel C. Hannah, Trudy F. C. Mackay, Robert R. H. Anholt

**Affiliations:** Department of Genetics and Biochemistry and Center for Human Genetics, Clemson University, Greenwood, SC29646, USA

**Keywords:** behavioral genetics, single cell RNA sequencing, transcriptomics, model organism, Fetal Alcohol Spectrum Disorder, interaction networks

## Abstract

Fetal alcohol exposure can lead to developmental abnormalities, intellectual disability, and behavioral changes, collectively termed fetal alcohol spectrum disorder (FASD). In 2015, the Centers for Disease Control found that 1 in 10 pregnant women report alcohol use and more than 3 million women in the USA are at risk of exposing their developing baby to alcohol. *Drosophila melanogaster* is an excellent genetic model to study developmental effects of alcohol exposure because many individuals of the same genotype can be reared rapidly and economically under controlled environmental conditions. Flies exposed to alcohol undergo physiological and behavioral changes that resemble human alcohol-related phenotypes. Here, we show that adult flies that developed on ethanol-supplemented medium have decreased viability, reduced sensitivity to ethanol, and disrupted sleep and activity patterns. To assess the effects of exposure to alcohol during development on brain gene expression, we performed single cell RNA sequencing and resolved cell clusters with differentially expressed genes which represent distinct neuronal and glial populations. Differential gene expression showed extensive sexual dimorphism with little overlap between males and females. Gene expression differences following developmental alcohol exposure were similar to previously reported differential gene expression following cocaine consumption, suggesting that common neural substrates respond to both drugs. Genes associated with glutathione metabolism, lipid transport, glutamate and GABA metabolism, and vision feature in sexually dimorphic global multi-cluster interaction networks. Our results provide a blueprint for translational studies on alcohol-induced effects on gene expression in the brain that may contribute to or result from FASD in human populations.

## Introduction

Prenatal exposure to ethanol can trigger a wide range of adverse physiological, behavioral, and cognitive outcomes, collectively termed fetal alcohol spectrum disorder (FASD) (1–4). Fetal alcohol syndrome (FAS) has the most severe manifestations of all FASDs, including craniofacial dysmorphologies, neurocognitive deficiencies, and behavioral disorders such as hyperactivity, attention deficit disorder and motor coordination anomalies (1,5–7). FAS/FASD is the most common preventable pediatric disorder, often diagnostically confounded with autism spectrum disorder (8). Time, dose, and frequency of exposure are often unknown, and manifestations of FASD are diverse and become evident long after exposure. The Centers for Disease Control and Prevention found that 1 in 10 pregnant women report alcohol use and more than 3 million women in the USA are at risk of exposing their developing baby to alcohol, despite warning labels on alcoholic beverages that indicate possible effects on prenatal development (9). Adverse consequences of fetal alcohol exposure extend throughout the lifespan.

Determining the effects of developmental alcohol exposure on adult phenotypes and gene expression in the adult brain is challenging in human populations, but can be addressed in model organisms. *Drosophila melanogaster* is an excellent model to study developmental effects of alcohol exposure, as we can control the genetic background and environmental conditions for large numbers of individuals without regulatory restrictions and at low cost. Importantly, flies exposed to alcohol experience loss of postural control, sedation, and development of tolerance (10–13), resembling human alcohol intoxication. Previous studies on the effects of developmental alcohol exposure in Drosophila showed reduced viability and delayed development time (14,15), reduced adult body size (14) and disruption of neural development (16). Developmental exposure to alcohol was associated with reduction in the expression of a subset of insulin-like peptides and the insulin receptor (14), dysregulation of lipid metabolism and concomitant increased oxidative stress (17), and reduced larval food intake due to altered neuropeptide F signaling (18).

Here, we show that developmental alcohol exposure in Drosophila results in decreased viability, reduced sensitivity to ethanol and disrupted sleep and activity patterns. Single cell RNA sequencing on adult fly brains following developmental alcohol exposure shows widespread sexually dimorphic changes in gene expression. These changes in gene expression resemble changes observed previously following cocaine exposure (19), indicating common neuronal and glial elements that respond to alcohol and cocaine consumption.

## Materials and Methods

### Drosophila Stocks and Exposure to Ethanol

*D. melanogaster* of the wild type Canton S (B) strain were maintained on cornmeal/yeast/molasses-agar medium supplemented with yeast at 25°C on a 12h light:dark cycle with 50% humidity, in controlled adult density vials to prevent overcrowding. We allowed 5 males and 5 females to mate for two days and aged their progeny for 3-5 days after eclosion. We then placed 50 males and 50 females into large egg collection cages on grape juice agar and yeast paste. We acclimatized the flies to the cages for 24 hours with grape juice plate changes every 12 hours, and collected up to 12-hour old eggs with a blunt metal needle. We placed the eggs on cornmeal-agar-molasses medium (control) or on cornmeal-agar-molasses medium containing 10% (v/v) ethanol (ethanol) without yeast. We collected 50 eggs per vial and set up 10-15 vials per condition per collection week over a 48-hour period (Figure 1). After eclosion, flies were transferred to control medium without yeast and aged as indicated for the relevant experiments. Unless otherwise indicated, all behavioral assays were performed in a controlled environment at 25°C.

**Figure 1.**
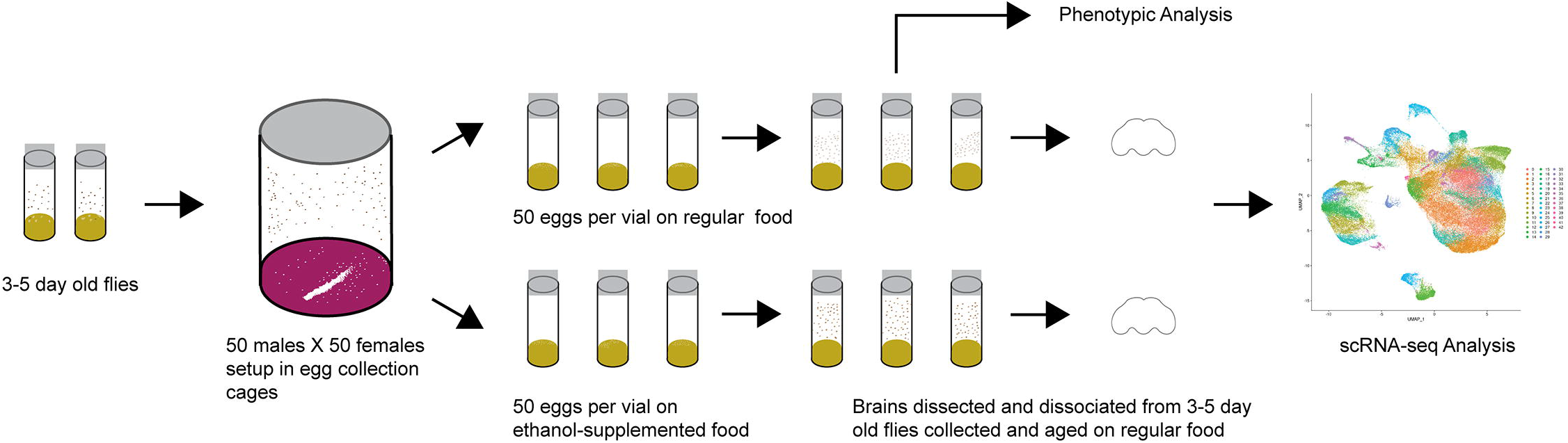
Diagram of the experimental design.

### Viability

The number of flies that emerged from vials into which 50 eggs had been placed were counted and the data were analyzed using the “PROC GLM” command (Type III) in SAS v3.8 (Cary, NC) according to the model *Y* = μ + *T* + ∊, where *Y* is the number of eclosed flies, μ is the population mean, *T* is the fixed effect of treatment (flies reared on control or ethanol medium), and ∊ is the residual error.

### Ethanol sensitivity

We measured ethanol sedation time as described previously (20) on 44-48 3-5 day old flies per sex per treatment. Ethanol sedation time was assessed between 8:30am and 11:30am. The number of seconds required for flies to lose postural control was analyzed using the “PROC GLM” command (Type III) in SAS v3.8 according to the model *Y* = μ + *T* + *S* + *TxS* + ∊, where *Y* is the time to sedation, μ is the population mean, *T* is the fixed effect of treatment (control or ethanol medium), *S* is the fixed effect of sex, *a*nd ∊ is the residual error.

### Sleep and Activity

Flies reared on either control or ethanol medium were placed in Drosophila Activity Monitors (DAM) (TriKinetics, Waltham, MA) containing a 5% sucrose, 2% agar medium at 1-2 days of age, and monitored for seven days on a 12 hour light-dark cycle. Activity was recorded as counts every time the fly interrupts an infrared beam. Sleep was defined as at least five minutes of inactivity. Only data from flies that survived the entire testing period were included, resulting in 57-64 flies per sex per treatment for analysis. Raw DAM monitor data were run in ShinyR-DAM (21), and the outputs were downloaded and parsed according to phenotype (e.g. day/night, sleep/activity, bout length/bout count) for subsequent statistical analyses. The data were analyzed using the “PROC MIXED” command (Type III) in SAS v3.8 according to the model *Y* = μ + *T* + *S* + *TxS* + *Rep(TxS)* + ∊, where *Y* is the sleep or activity phenotype, μ is the population mean, *T* is the fixed effect of treatment (control or ethanol medium), *S* is the fixed effect of sex, *Rep* is the random effect of replicate and ∊ is the residual error. Reduced models were also performed for each sex.

### Brain Dissociation and Single Cell RNA Sequencing

For single cell RNA sequencing, we collected duplicate samples of 20 brains for each sex from flies reared on control or ethanol medium. We dissociated the brains as previously described after incubation with 450μl of collagenase solution (50 ul of fresh 25mg/ml collagenase (Gibco) in sterile water + 400μl of Schneider’s medium) for 30 minutes followed by stepwise trituration - P200 pipette 5 times, 23G needle pre-wetted with PBS + BSA 5 times, and 27G pre-wetted needle 5 times (19). The resulting suspension was passed through a pre-wetted 10μm strainer (Celltrics, Görlitz, Germany) with gentle tapping. We counted live cells using a hemocytometer with trypan blue exclusion and proceeded with GEM generation using the Chromium controller (10X Genomics, Pleasanton, CA) for samples with > 500 live cells/μl. We prepared libraries in accordance with 10X Genomics v3.1 protocols. We determined fragment sizes using Agilent Tapestation kits (Agilent, Santa Clara, CA) - d5000 for amplified cDNA and d1000 for libraries. We measured the concentrations of amplified cDNA and final libraries using a Qubit 1X dsDNA HS kit (Invitrogen, Waltham, MA) and a qPCR based library quantification kit (KAPA Biosystems, Roche, Basel, Switzerland). We used 12 cycles for the cDNA amplification and 12 cycles for indexing PCR. We sequenced the final libraries on an Illumina NovaSeq6000.

### Single Cell RNA Sequencing Data Analysis and Bioinformatics

We used the *mkfastq* pipeline within Cell Ranger v3.1 (10X Genomics, Pleasanton, CA) to convert BCL files from the sequence run folder to demultiplexed FASTQ files. We used the *mkref* pipeline to index the release 6 version of the *D. melanogaster* reference *GCA_000001215.4* from NCBI Genbank. For alignment, we used the *count* pipeline within Cell Ranger v3.1 with the expected cell count parameter set to 5,000 cells. We imported raw expression counts output for each sample from the Cell Ranger pipeline and analyzed these data using the Seurat v3 package in R (22). We normalized counts by regularized negative binomial regression using the *scTransform* pipeline (23). We performed integration of samples using the *SCT* method. *RunUMAP* and *FindNeighbors* functions were used with 10 dimensions to ordinate expression space and reduce data dimensionality. To identify cell-type clusters, we used unsupervised clustering using the *FindClusters* function and assigned the origin of clustered cells based on well-established biomarkers.

We used the Pearson residuals output from the *scTransform* pipeline as input for differential expression calculation (23). We used the *MAST* algorithm as the testing methodology in the *FindMarkers* function for each cluster to calculate differential expression, which allows for the incorporation of the cellular detection rate, defined as a fraction of genes expressed in each cell, as a covariate (24). *P-*values for differential expression were adjusted for multiple-hypothesis testing using a Bonferroni correction, and adjusted *p*-values that are less than 0.05 were considered statistically significant.

Interaction networks were produced using the unique list of differentially expressed genes aggregated from all clusters and the stringApp (25) within Cytoscape (26).

The code for all analyses can be found here: https://github.com/vshanka23/The-Drosophila-Brain-after-developmental-ethanol-exposure-at-Single-Cell-Resolution/blob/main/Rcode_for_analysis.R

## Results

### Effects of Developmental Alcohol Exposure on Adult Phenotypes

Exposure of flies to ethanol during the embryonic and larval stages resulted in an 8.9% reduction in viability compared to flies reared on control medium (Figure 2A). The adult flies exposed to ethanol during development did not show any overt morphological abnormalities. We next asked whether developmental alcohol exposure would alter sensitivity to acute alcohol exposure as adults. We reared developing flies on ethanol medium and transferred the adults to control medium immediately after eclosion. The flies that developed on ethanol medium showed reduced sensitivity (longer sedation times) to acute alcohol exposure in both sexes, indicating increased tolerance to acute alcohol exposure compared to flies that developed on control medium (Figure 2B).

**Figure 2.**
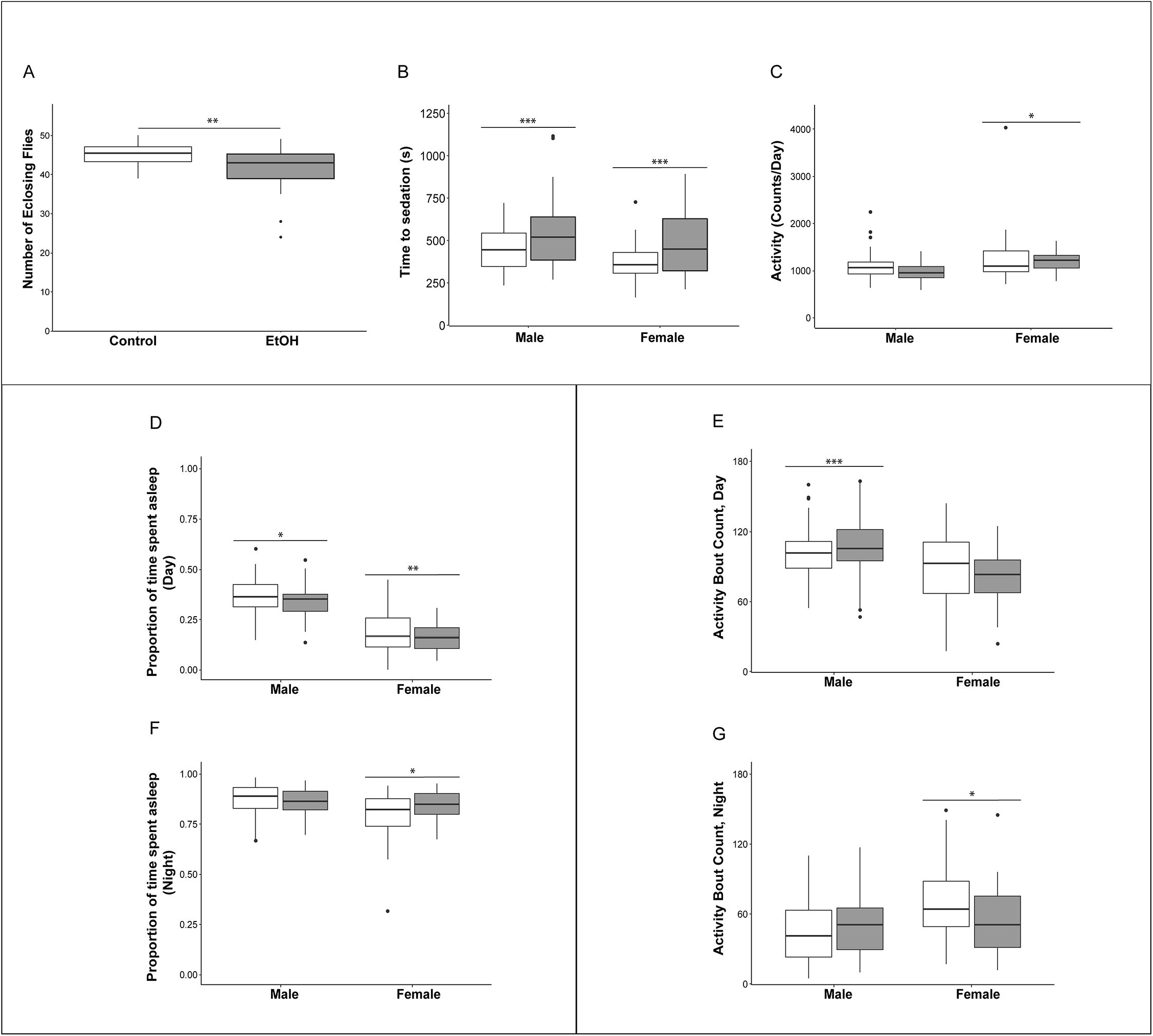
Effects of developmental alcohol exposure on viability and behavioral phenotypes in adult flies. (A) Boxplots of viability (n=12 reps of 50 embryos per treatment), (B) Ethanol sensitivity (n=43-49 3-5 day old flies per sex per treatment), (C) Activity, (D) Proportion of daytime sleep, (E) Activity bouts during the day, (F) Proportion of night time sleep, (G) Activity bouts during the night. Day hours are from 6am-6pm. Grey boxes indicate flies reared on medium supplemented with 10% (v/v) ethanol and white boxes indicate control flies grown on regular medium. n=57-64 flies per sex per treatment for all sleep and activity phenotypes. * *p*<0.05, ** *p*<0.01, *** *p*<0.001.

Children with FASD often have disturbed sleep (27, 28). Therefore, we used the Drosophila Activity Monitor system to assess the effects of developmental alcohol exposure on adult activity and sleep patterns, and found that exposure to alcohol during development had sex-specific effects on these phenotypes. Overall activity in males was not affected by the ethanol treatment, but females exposed to ethanol were more active (Figure 2C; Supplementary Table S1). Ethanol exposure reduced sleep during the day in both sexes (Figure 2D), and day sleep in males was fragmented, with an increase in activity bouts (Figure 2E). In contrast, females compensated for increased activity and reduced daytime sleep with extended periods of night sleep (Figure 2F) with a reduced number of activity bouts (Figure 2G; Supplementary Table S1).

### Effects of Developmental Alcohol Exposure on Gene Expression in the Brain

We performed single cell RNA sequencing to assess the effects of developmental alcohol exposure on gene expression in the brain in males and females, with two replicates per sex and treatment (Figure 1). We obtained a total of 108,571 cells across all samples, which corresponds to ~10% of all cells in a Drosophila brain (Supplementary Table S2). We visualized these data using the Uniform Manifold Approximation and Projection (UMAP) non-linear dimensionality reduction method (29), which showed that all samples were uniformly represented (Figure 3; Supplementary Table S2). Unsupervised clustering of the dataset generated 43 cell clusters, which represent the major regions of the Drosophila brain, including neuronal and glial populations, and all major neurotransmitter cell types (Figure 4; Supplementary Table S3). We identified seven distinct populations of GABAergic neurons, two subpopulations of Kenyon cells of the mushroom bodies (integrative centers for experience-dependent modulation of behavior), and several distinct populations of glia, including two separate clusters of astrocytes as well as surface glia that form the blood-brain barrier (Figure 4).

**Figure 3.**
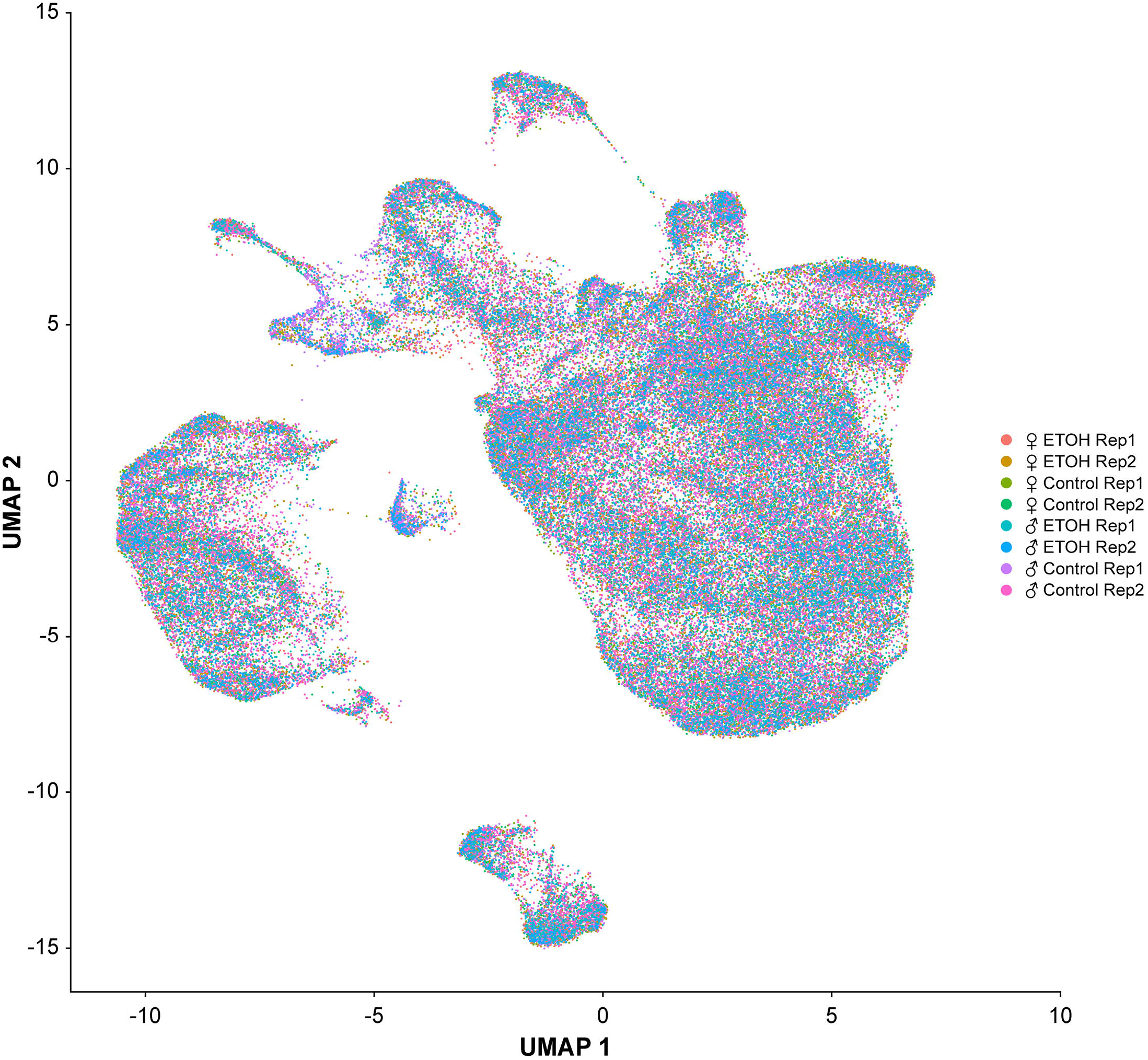
Uniformity across samples of single cell transcriptomes. Gene expression patterns of single cells (n = 108,571) from all eight samples are represented in low dimensional space using a graph-based, non-linear dimensionality reduction method (UMAP). Individual dots represent the transcriptome of each cell and the colors of the dots represent the samples to which the cells belong.

**Figure 4.**
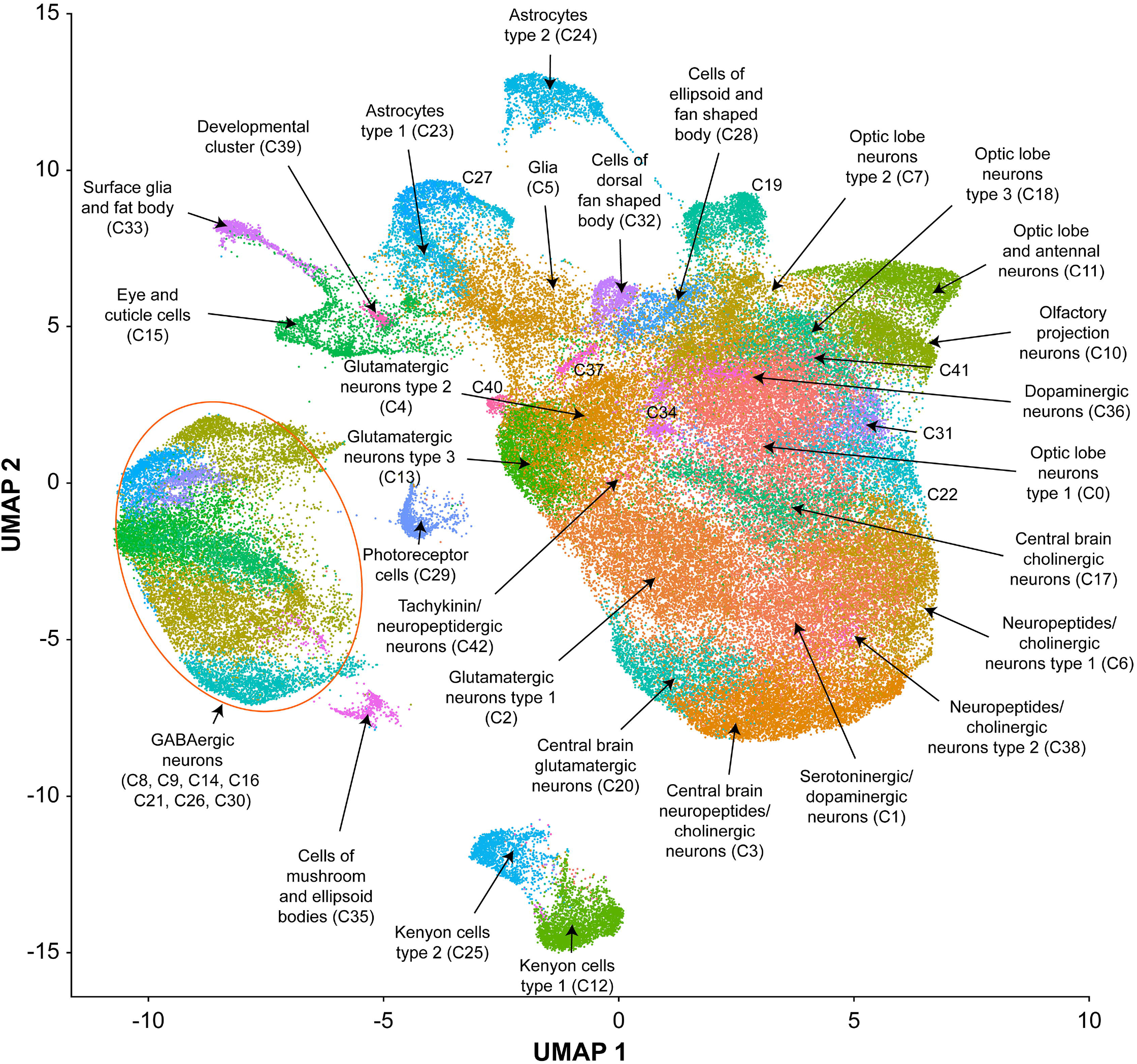
UMAP visualization and annotation of cell clusters. Cells were clustered based on their expression pattern using the unsupervised shared nearest neighbor (SNN) clustering algorithm. Individual dots represent each cell and the colors of the dots represent the cluster to which the cells belong. Annotation of cell types from clusters was performed by cross-referencing cluster-defining genes across FlyBase (50) and published literature (Supplementary Table S3).

We combined all differentially expressed genes from all clusters and performed differential expression analyses. We found 119 transcripts in males and 148 transcripts in females with altered abundances after developmental alcohol exposure at a Bonferroni adjusted *p*-value <0.05. We identified 61 upregulated and 25 downregulated genes in males, and 57 upregulated and 34 downregulated genes in females at a threshold of |log_e_FC| > 0.25 (Figure 5; Supplementary Tables S4 and S5). Increasing the stringency to |log_e_FC| > 1.0 (Bonferroni adjusted *p* value <0.05) retained 36 upregulated and 10 downregulated genes in males and 32 upregulated and 20 downregulated genes in females (Supplementary Figure S1). Differential expression patterns are sexually dimorphic, as observed previously for cocaine-induced modulation of gene expression (19), with only 32 differentially expressed genes in common between the sexes. Changes in gene expression in the mushroom bodies, represented by cluster C12, are primarily observed in females. Developmental alcohol exposure modulates expression of several genes in glia, represented by clusters C5, C15, C23, C24, and C33, in a sexually dimorphic pattern (Figure 5). Especially noteworthy is the prominent differential expression of *lncRNA:CR31451*, a long non-coding RNA of unknown function, in multiple neuronal populations. This transcript is globally upregulated in males but downregulated in females (Figure 5; Supplementary Figure S1). Among all differentially expressed genes, ~ 58% have human orthologs (DIOPT score ≥ 3; Supplementary Table S6).

**Figure 5.**
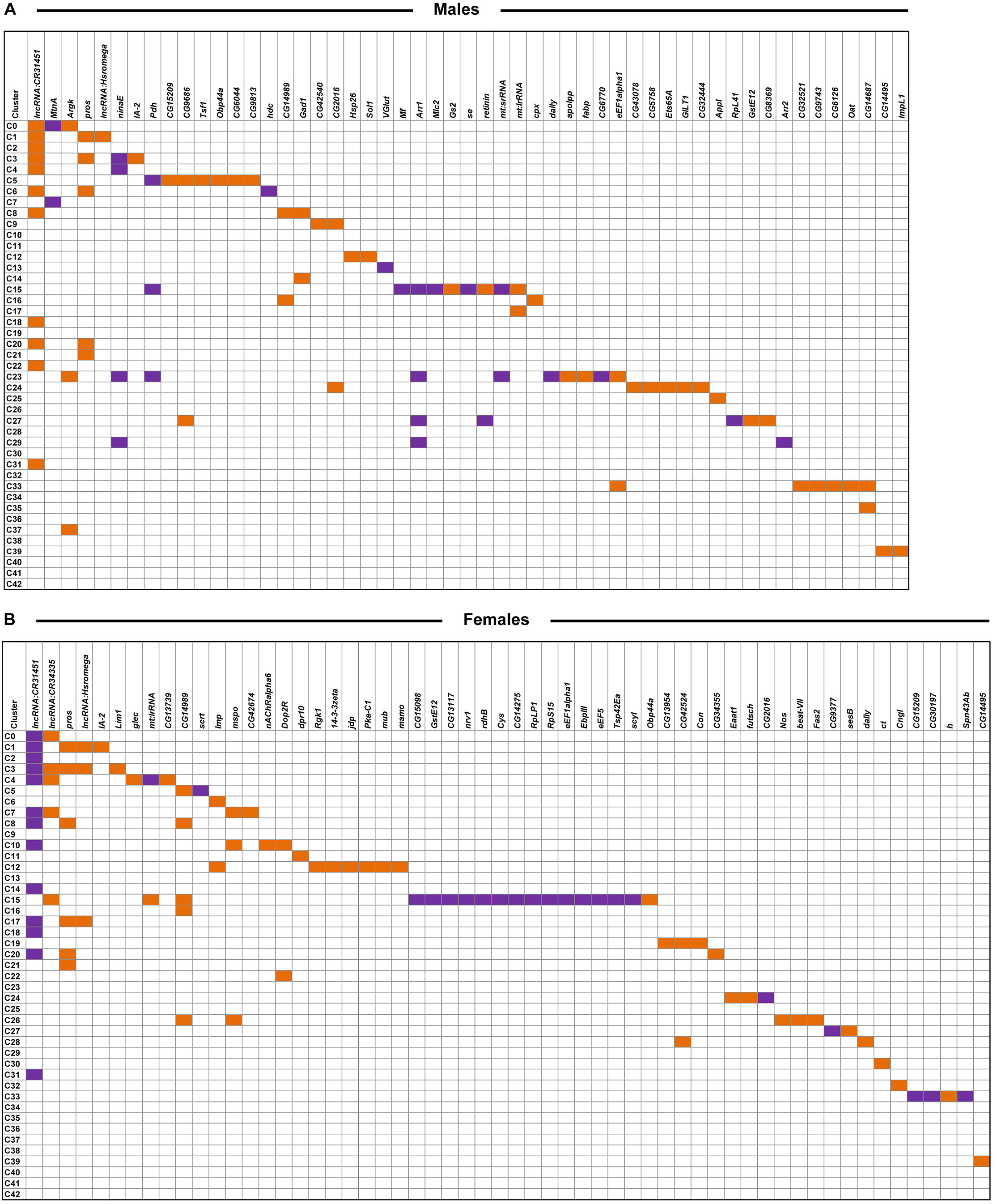
Differentially expressed genes across clusters in males (A) and females (B) after developmental alcohol exposure. Differentially expressed genes are listed on the top (columns) and cell clusters are represented by the rows. Upregulated genes are indicated with orange and downregulated genes are indicated with purple. Differentially expressed genes are filtered at |log_e_FC| > 0.25 and a Bonferroni adjusted *p* value <0.05. Differentially expressed genes that survive a threshold of |log_e_FC| > 1.0 with a Bonferroni adjusted *p* value <0.05 are shown in Supplementary Figure S1.

We assessed global interaction networks of differentially expressed gene products across all cell clusters for males and females separately (Figure 6). The male interaction network is composed of modules associated with glutathione metabolism, lipid transport, glutamate and GABA metabolism, and vision (Figure 6A). The female interaction network also contains modules associated with glutamate and GABA metabolism, lipid metabolism, and vision, but the composition of these modules is distinct from their male counterparts. In addition, the female network features modules associated with monoaminergic signaling, cell adhesion, and Wnt signaling (Figure 6B). Multiple cell clusters contribute to each network module, indicating that modulation of gene regulation by developmental alcohol exposure is coordinated across different cells throughout the brain.

**Figure 6.**
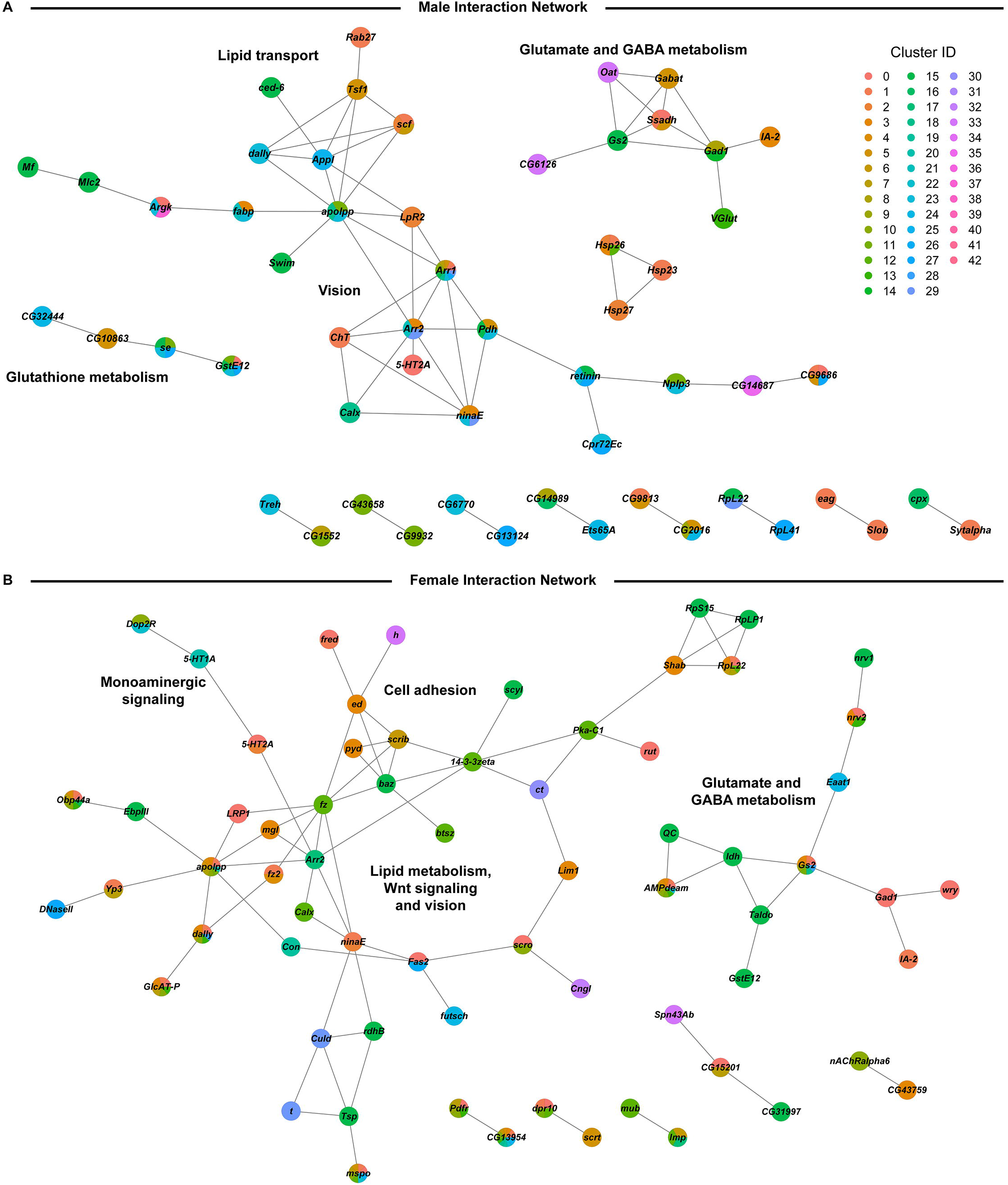
Global interaction networks of differentially expressed gene products in males (A) and females (B) following developmental alcohol exposure. Colors of the nodes correspond to the clusters in which expression of the gene is altered after growth on alcohol-supplemented medium.

We noticed that many genes that are differentially expressed following developmental exposure to ethanol correspond to genes that undergo altered expression when flies are exposed to cocaine (19). However, the transcriptional response to acute exposure to cocaine is larger than the transcriptional response to developmental alcohol exposure. Nonetheless, 69.7% of differentially expressed genes in males and 43.2% of differentially expressed genes in females in our data overlap with differentially expressed genes after consumption of cocaine (Figure 7; Supplementary Table S7), although the magnitude and direction of differential expression of common genes between the two treatments varies by cell type (Supplementary Table S8). Gene ontology enrichment analyses of this common set of genes in each sex highlights gene ontology categories associated with development and function of the nervous system (Supplementary Table S9, 30).

**Figure 7.**
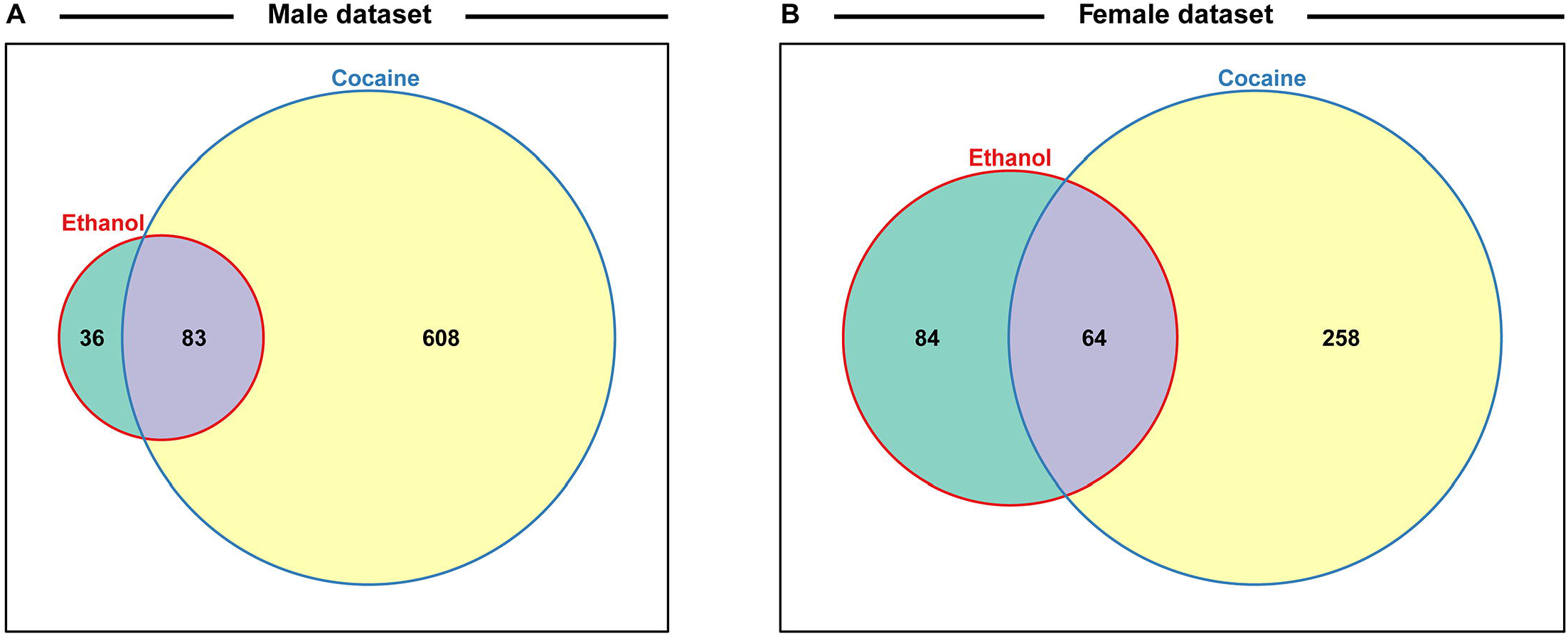
Venn diagrams indicating the proportions of differentially regulated genes after exposure to alcohol during development or acute consumption of cocaine for males (A) and females (B). Data for cocaine exposure are from ref 19. See also Supplementary Table 7.

## Discussion

We characterized the consequences of developmental alcohol exposure in Drosophila on viability, behavioral phenotypes, and gene expression in the brain. Characteristic features of FASD in humans include craniofacial dysmorphologies and cognitive impairments. Although we did not perform detailed morphometric measurements, we did not observe any overt morphological aberrations, and cognitive impairments are challenging to assess in Drosophila. Nevertheless, flies exposed to alcohol during embryonic and larval development showed changes in activity and sleep patterns (Figure 2C-G), reminiscent of activity and sleep disturbances seen in children with FASD (27, 28). We also find that growth on alcohol supplemented medium results in reduced ethanol sensitivity of adult flies, in agreement with a previous study (Figure 2B, 14).

We hypothesize that the effects of developmental alcohol exposure on changes in gene expression in the Drosophila central nervous system will converge on evolutionarily conserved cellular processes. Drosophila is advantageous for studies on gene expression at single cell resolution because we can survey the entire brain in a single analysis, unlike studies in rodents, and pooling multiple brains of the same genotype averages individual variation. The power to detect changes in gene expression in our study is improved by only considering changes in gene expression that are consistent across replicates.

We observed changes in gene expression in adult flies, even though exposure to alcohol occurred only during the larval stages and briefly after eclosion, after which adults were collected and maintained on regular medium without alcohol. It is possible that developmental alcohol exposure may result in epigenetic modifications that give rise to altered gene expression patterns into adulthood (31).

We observe changes in gene expression in diverse neuronal and glial cell populations (Figure 5). Since we are not able to sample all cells of the brain, it is likely that some neuronal or glial cell populations are not represented in our data. However, the major regions of the Drosophila brain and all major neurotransmitter cell types are represented (Figure 4; Supplementary Table S3). The effects of developmental alcohol exposure are sexually dimorphic, similar to previously observed changes in transcript abundances following consumption of cocaine (19). Sexual dimorphism is also a hallmark of FASD, with different effects of fetal alcohol exposure on neural development and cognitive abilities between males and females (32–35). Although different genes are affected in males and females, gene ontology analysis indicates that they converge on the same biological processes, related to development and function of the nervous system (Table S8). The considerable overlap between differentially expressed genes in response to alcohol and cocaine suggests common neural substrates that respond to toxic exposures. Genes associated with immune defense and xenobiotic detoxification, including the glutathione pathway, feature in interaction networks of differentially expressed gene products (Figure 6).

*lncRNA:CR31451* shows large sexually antagonistic responses to developmental alcohol exposure in many neuronal cell populations. Whereas a previous study documented expression of this gene in glia (36), we only observe differential gene expression of *lncRNA:CR31451* in neurons under the conditions of our study (Figure 5). Future studies are needed to assess whether this gene product fulfills a regulatory function that affects multiple neurotransmitter signaling processes and whether its sex-antagonistic response to alcohol exposure could in part cause the differential gene expression patterns seen in males and females.

Our observations of extensive changes in gene expression in glia in response to developmental alcohol exposure are in accordance with the role of glia in FASD. Fetal alcohol exposure leads to impaired astrocyte development and differentiation, which gives rise to microencephaly (37, 38). In addition, ethanol exposure increases permeability of the blood brain barrier (39), which in Drosophila is formed by the surface glia (40). Among the glial genes that show altered expression after developmental alcohol exposure in Drosophila are *GILT1*, which contributes to the immune defense response to bacteria (41), *Gs2* and *Eaat1*, which are involved in glutamine synthesis and transport of glutamate in astrocytes (42, 43), *GstE12* and *se*, which are involved in glutathione metabolism (44), and *fabp* and *apolpp*, which function in lipid metabolism (45, 46).

GABA signaling and glutamate signaling neuronal cell populations feature prominently in our data (Figure 3). Glutamate is also a precursor for the biosynthesis of glutathione, which is produced in glia and protects against oxidative stress and detoxification of xenobiotics (47). Developmental alcohol exposure interferes with glutamate and GABA signaling because ethanol is both an antagonist to the NMDA glutamate receptor and mimics GABA (48). Consequently, fetal alcohol exposure results in neuronal apoptosis during the rapid brain growth spurt during which the astrocytes play a major role (48, 49). Evolutionarily conserved neural processes that respond to developmental alcohol exposure in Drosophila thus provide a blueprint for translational studies on alcohol-induced effects on gene expression in the brain that may contribute to or result from FASD in human populations.

## Data Availability Statement

The datasets for this study can be found in the GEO repository under accession number GSE172231.

## Author Contributions

SSM and VS contributed equally; SSM and RCH maintained fly stocks, reared flies on developmental alcohol, and measured viability; RAM measured and analyzed ethanol sensitivity and sleep and activity phenotypes; SSM and RCH performed brain dissociation; SSM performed RNA sequencing; SSM and VS analyzed the RNA sequencing data; SSM, TFCM and RRHA conceived of the experiments; SSM, VS, RAM, TFCM and RRHA wrote the manuscript; TFCM and RRHA provided resources.

## Supplementary Materials

**Supplementary Figure S1. Differentially expressed genes across clusters in males (A) and females (B) after developmental alcohol exposure.** Differentially expressed genes are listed on the top (columns) and cell clusters are represented by the rows. Upregulated genes are indicated with orange and downregulated genes are indicated with purple. Differentially expressed genes are filtered at |log_e_FC| > 1.0 and a Bonferroni adjusted *p* value <0.05.

**Supplementary Table S1. ANOVA tables for viability, ethanol sensitivity, and sleep and activity.**

**Supplementary Table S2. Sequencing statistics.** F denotes females and M denotes males. C indicates control medium and E ethanol-supplemented medium. The numbers indicate replicates 1 and 2.

**Supplementary Table S3. Genes used to annotate cell clusters.**

**Supplementary Table S4. List of differentially expressed genes in each cluster in males.** Each sheet corresponds to the male analyses for the given cluster. “Avg_diff” is conditionally formatted to indicate up- and down-regulation of expression in ethanol compared to regular food (red: up-regulated, green: down-regulated and yellow: no difference). p_val: raw p-value from the differential expression analysis for the given gene in the corresponding cluster. avg_diff: the difference in the log(e) transformed average expression of the given gene in the corresponding cluster (sheet) between the two conditions (ethanol compared to regular food). Values above zero indicate up-regulation of expression due to developmental exposure to ethanol, and likewise, values below zero represent down-regulation of expression due to ethanol. p_val_adj: Bonferroni adjusted p-value. The DE matrix sheet is a summary of differentially expressed genes (columns) and the clusters in which they are differentially expressed (rows) with orange indicating upregulation and purple indicating downregulation at |avg_diff| thresholds of 0.25 and 1. The All DE per cluster sheet and the All DE sheet are summaries of all the differentially expressed genes.

**Supplementary Table S5. List of differentially expressed genes in each cluster in females.** Each sheet corresponds to the female analyses for the given cluster. “Avg_diff” is conditionally formatted to indicate up- and down-regulation of expression in ethanol compared to regular food (red: up-regulated, green: down-regulated and yellow: no difference). p_val: raw p-value from the differential expression analysis for the given gene in the corresponding cluster. avg_diff: the difference in the log(e) transformed average expression of the given gene in the corresponding cluster (sheet) between the two conditions (ethanol compared to regular food). Values above zero indicate up-regulation of expression due to developmental exposure to ethanol, and likewise, values below zero represent down-regulation of expression due to ethanol. p_val_adj: Bonferroni adjusted p-value. The DE matrix sheet is a summary of differentially expressed genes (columns) and the clusters in which they are differentially expressed (rows) with orange indicating upregulation and purple indicating downregulation at |avg_diff| thresholds of 0.25 and 1. The All DE per cluster sheet and the All DE sheet are summaries of all the differentially expressed genes.

**Supplementary Table S6. Human orthologs of differentially expressed genes.**

**Supplementary Table S7. Common differentially expressed genes upon developmental alcohol exposure and acute exposure to cocaine.**

**Supplementary Table S8. Comparison of cell type-specific differentially expressed genes between developmental ethanol exposure and acute cocaine exposure.** Meta-comparison sheet contains the mapping of clusters and cell types between the two datasets as well as the methodology and summary of the comparisons. The rest of the sheets contain the list of statistically significantly differentially expressed genes, their Log_e_ fold change values, the calculations of the comparisons between the two datasets for each cell type-category. The comparisons were done for each cell type-category separately for the male and female datasets.

**Supplementary Table S9. Gene ontology analysis of differentially expressed genes identified both after developmental exposure to alcohol and acute intake of cocaine.**

