## Supplementary figures and images for "Developmental Alcohol Exposure in Drosophila: Effects on Adult Phenotypes and Gene Expression in the Brain"

### Supplementary Figure S1

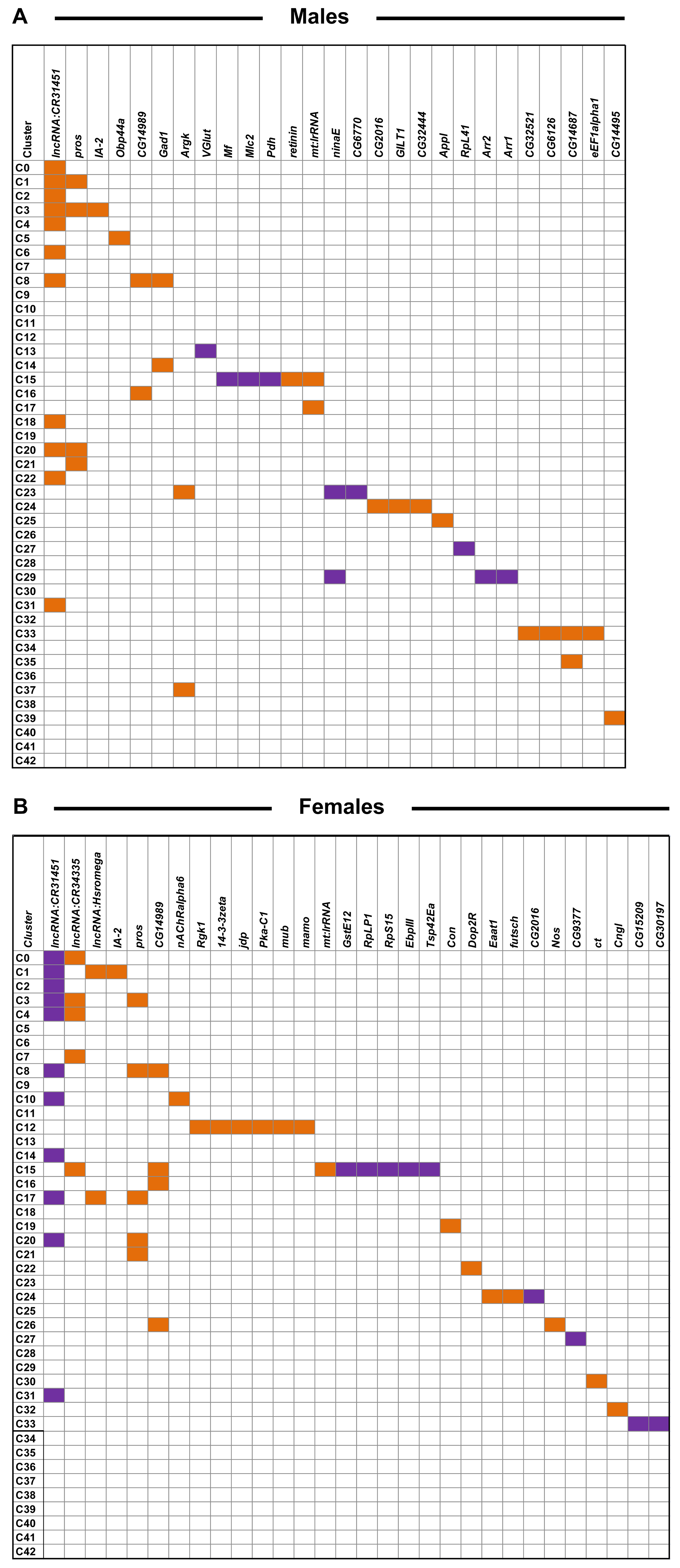
